# Cell-Type Clusters in the MICrONS Connectome Reveal Hidden Organizational Principles of the Mouse Visual Cortex and Possible Candidate Substrates for Elementary Perceptual States

**DOI:** 10.64898/2026.05.12.724707

**Authors:** W. Alexander Escobar

## Abstract

How neural structure constrains cortical dynamics and perceptual experience remains a fundamental open question. The MICrONS connectomics dataset enables direct examination of this problem by combining dense ultrastructural reconstructions with functional measurements across a mesoscale volume of mouse visual cortex. Here, I test a prediction arising from field-based models of perception: that subsets of cortical neurons should exhibit rare but reproducible structural similarity and spatial clustering.

Using exhaustive pairwise NBLAST comparisons across tens of thousands of reconstructed neurons, I quantified morphological similarity among excitatory populations spanning layers 2/3, 5, and 6, including extratelencephalic, intratelencephalic, and near-projecting cells. High similarity scores were exceedingly rare, occurring more than 8–10 standard deviations above population means. Despite this rarity, structurally similar neurons formed discrete, spatially coherent clusters across primary and higher-order visual areas, often aligning across cortical layers. These findings reveal a previously unrecognized level of mesoscale structural organization and provide an empirical foundation for linking cortical microarchitecture to field-based models of visual experience.

## Introduction

Understanding how neural structure constrains neural function remains a central challenge for systems neuroscience. Recent advances in large-scale connectomics have begun to bridge this gap by enabling the simultaneous examination of cellular morphology, synaptic connectivity, and functional responses within the same tissue. Among these resources, the MICrONS Connectomics Dataset provides an unprecedented view of neural circuitry across a mesoscale cortical volume (1.4 × 0.87 × 0.84 mm) of mouse visual cortex (The MICrONS Consortium et al. 2025); (Turner et al. 2022). This dataset contains dense reconstructions of tens of thousands of neurons spanning all six cortical layers, together with in vivo two-photon calcium activity collected during visual stimulation. Such multimodal richness permits investigation of how fine-scale dendritic architecture, connectivity, and physiological responses jointly shape cortical computation.

Neurons of the primary visual cortex form dense, recurrent networks through extensive local and long-range synaptic connectivity. Such recurrent architectures are known to support attractor dynamics, in which population activity evolves toward stable or quasi-stable states within a high-dimensional activity space. These attractor states reflect reproducible patterns of excitation and inhibition across the network and have been widely proposed as neural substrates for learning, memory, and internal models of sensory context (Hopfield 1982); (Amit and Brunel 1997); (Yang et al. 2019).

When a network settles into a particular attractor state, it generates a structured, repeatable pattern of synaptic drive onto target neurons. For a given postsynaptic neuron, this attractor-specific balance of excitation and inhibition sculpts a characteristic distribution of transmembrane currents across its dendritic arbor. These currents, in turn, give rise to a spatially organized electromagnetic field whose topology reflects both the neuron’s morphology and the attractor-imposed pattern of synaptic activation. Because attractor states are stable yet revisitable, the resulting electromagnetic field configurations are expected to be metastable—persisting for behaviorally relevant timescales while remaining dynamically accessible—thus providing a potential physical instantiation of circuit-level internal models at the level of single-neuron field dynamics.

Herms Romijn proposed in 2002 that “Based on neurobiological data, modern concepts of self-organization and a careful rationale, the hypothesis is put forward that the fleeting, highly ordered patterns of electric and/or magnetic fields, generated by assemblies of dendritic trees of specialized neuronal networks, should be thought of as the end-product of chaotic, dynamically governed self-organization” (Romijn 2002). Romijn believed these dendritic arbor EM fields serve as the physical surrogates of consciousness, creating bits or quanta of conscious experience like a point of the color blue.

Importantly, this hypothesis does not require exact invariance in arbor topology across instances of a given stereotyped perceptual experience. Rather, it posits that dendritic morphologies constrained within a bounded physical regime may support the same effective electromagnetic field (EMF) topology, provided that patterns of synaptic excitation and inhibition are appropriately tuned. The Quantized Visual Awareness (QVA) hypothesis predicts the existence of neuronal classes whose gross structural features are sufficiently conserved to define discrete categories, each corresponding to a specific type of conscious visual experience (Escobar, 2011, 2013, 2016).

Moreover, QVA predicts that such structurally related neurons, together with the local circuits that drive them, are not randomly distributed within primary visual cortex but instead exhibit spatial clustering. These clusters are expected to align with known functional architectures, including orientation-selective pinwheel domains (Crair et al. 1997) and chromatic “blob” regions (Bartfeld and Grinvald 1992) observed in primate V1.

Layer 5 extratelencephalic-projecting (ET) neurons are excellent candidates for producing these EMFs. These cells have significant and complex dendritic arborizations and are considered an endpoint for recurrent activation (Moberg and Takahashi 2022). In addition, ET cells of layer 5 have a significantly higher number of synapses compared to all other cells throughout the mouse primary visual cortex (Schneider-Mizell et al. 2025). Moreover, they are surrounded by intratelencephalic (IT) cells that project onto ET and other cells forming highly complex neural networks. These neural networks might allow for the specific activation of ET cells required to form the appropriate EMF that serves as a quantum of experience.

To investigate whether morphological similarity indeed defines discrete subpopulations within the mouse visual cortex, I conducted an exhaustive pairwise NBLAST analysis across tens of thousands of neurons in the MICrONS volume. Consistent with theoretical predictions, I identified neuronal groups with pronounced morphological similarity that formed spatially coherent clusters within the cortical volume. These clusters reveal organizational principles not apparent from traditional cell-type classifications and suggest that structural similarity may play a previously unrecognized role in constraining the functional or dynamical properties of cortical circuits.

## Materials and Methods

### Comparing Neural Cells Using NBLAST

Each neuron to be compared was represented by a skeletonized 3D reconstruction, stored in the standard SWC format. These files contain the coordinates of neuronal points (nodes) and their connectivity, effectively describing the branching architecture of axons and dendrites. The skeletons for tens of thousands of neurons were downloaded from the MICrONS dataset using Google Colabs. SWC files are economical in terms of storage and given the large number of cells, it allowed for the rapid transfer of files to Google Drive and from there to a local server.

Before performing NBLAST, the neurons were imported into a Python environment capable of handling neuroanatomical data. I used the Navis library, which provides tools for reading, manipulating, and analyzing neuron skeletons.

All SWC files to be compared were placed in a single folder alongside the NBLAST comparison script (swc_nblast_similarity2.py). The script automatically detected all available SWC files, loaded them as TreeNeuron objects, and ensured that each reconstruction was standardized—centered on the soma or aligned to a consistent anatomical landmark. This alignment ensured that neurons were compared in the same spatial frame, reducing artifactual differences due to translation or rotation.

NBLAST does not compare raw skeletons directly. Instead, it relies on a dotprop representation, in which each node of a neuron is represented by a vector describing its spatial position and local tangent direction. This representation captures both the geometry and local orientation of neuronal processes. In Navis, this transformation is performed using the make_dotprops function, which converts TreeNeuron objects into Dotprops. Each Dotprop consists of: the 3D coordinates of sampled points along the neuron, a vector (the “dot”) representing the local direction of the neurite, and a uniform resampling step to ensure comparable density across neurons.

In addition, I used Navis’s nblast_smart function, which automatically selects the optimal computation strategy depending on the dataset size. It supports parallel processing, allowing simultaneous comparison across multiple cores, which greatly accelerates analysis for datasets containing hundreds or thousands of neurons.

After all pairwise comparisons were computed, the results were assembled into a similarity matrix, in which each cell (i, j) represents the NBLAST score between neuron *i* and neuron *j*. Neurons with higher pairwise scores are likely to share common morphological features—such as similar branching patterns, axon trajectories, or arborization territories—suggesting that they may belong to the same cell type or functional class.

### Grouping Cells based on their Score

To extract biologically meaningful relationships, neurons were grouped based on their NBLAST similarity scores. In this study, neurons that shared pairwise similarities greater than 0 were considered members of the same morphological group. This threshold empirically corresponds to neurons whose structures are visually and quantitatively comparable.

The grouping was performed computationally by examining all pairwise scores and linking neurons that exceed the similarity threshold. The resulting groups were listed in an output report, typically as “Group 1,” “Group 2,” and so forth, each containing the identifiers (the pt_root_ID from the MICrONS dataset) of its constituent neurons. This output was used as a foundation for downstream analysis, such as confirming morphological classes.

Due to the very large files (i.e. ∼19000×19000 ≈ 361 million .csv cells), loading the whole file at once was too heavy for memory or normal execution limits. Since it’s an upper-triangular similarity matrix, I exploited that structure to process it efficiently without ever holding the entire matrix in memory. The script (CELL_Nblast_Comparison.py) processed the file in chunks and output the RTF grouping file without loading the entire matrix into memory at once. CSV files are more memory-efficient and support chunked reading.

Multi-member groups are real clusters where each member is connected (directly or transitively) via edges > 0. If two nodes A and B are linked A–C–B via edges >0, they will be in same connected component (group) even if A–B direct score ≤0. This is expected behavior for connected components clustering.

## Results

Typical histograms taken from the population of each indicated cell type. All of these histograms represent scores for thousands of excitatory neurons within the mouse visual cortex.

Coordinates within the MICrONS dataset are generally given in voxels. To convert coordinate values into nanometers you must use the Standard Transform (https://tutorial.microns-explorer.org/quickstart_notebooks/08-standard-transform.html). Besides coverting the data to nanometers, this transform will flatten the pial service and set its depth (y coordinate) to zero. All the following representations of the volume are top-down views and contain no depth information.

## Discussion

The MICrONS dataset offers an unprecedented opportunity to examine the structural organization of mouse primary visual cortex (VISp) at synaptic resolution. Although the captured volume is anchored in VISp, it includes adjacent higher-order visual areas such as VISrl (rostrolateral) and VISal (anteriolateral), regions known to support disparity processing, orientation tuning, and contrast sensitivity (Figure 2A). This anatomical diversity situates the dataset within a functionally heterogeneous portion of the visual hierarchy, allowing structural analyses to be interpreted in the context of the well-established transformations that occur between early and downstream visual areas. The combination of dense EM reconstruction and large-scale cell-type annotations—including recent classifiers built on perisomatic features—further enables systematic comparisons across excitatory populations, including extra-telencephalic (ET), intra-telencephalic (IT), and near-projecting (NP) cells spanning all cortical layers (Elabbady et al. 2025).

**Figure 1.**
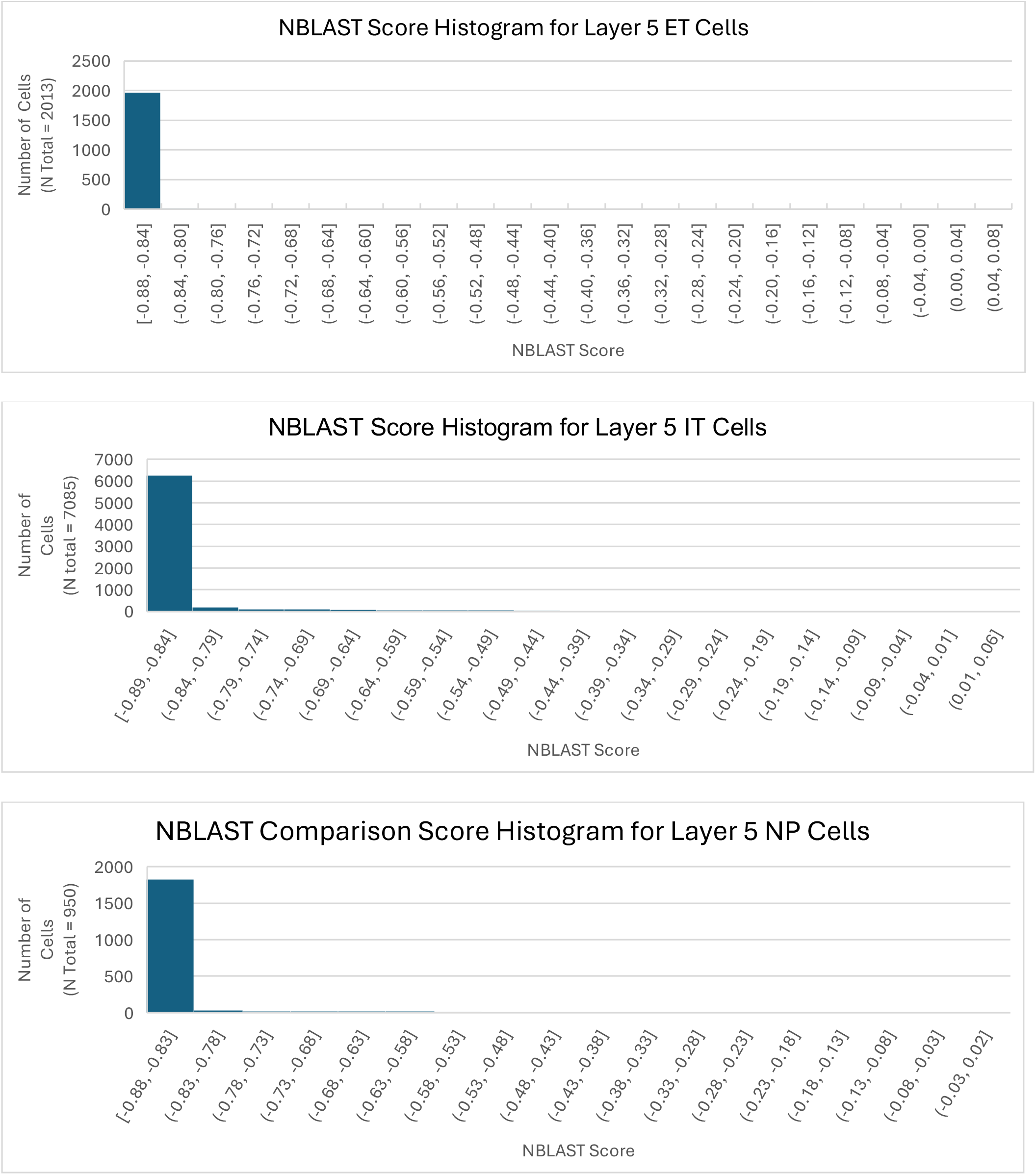
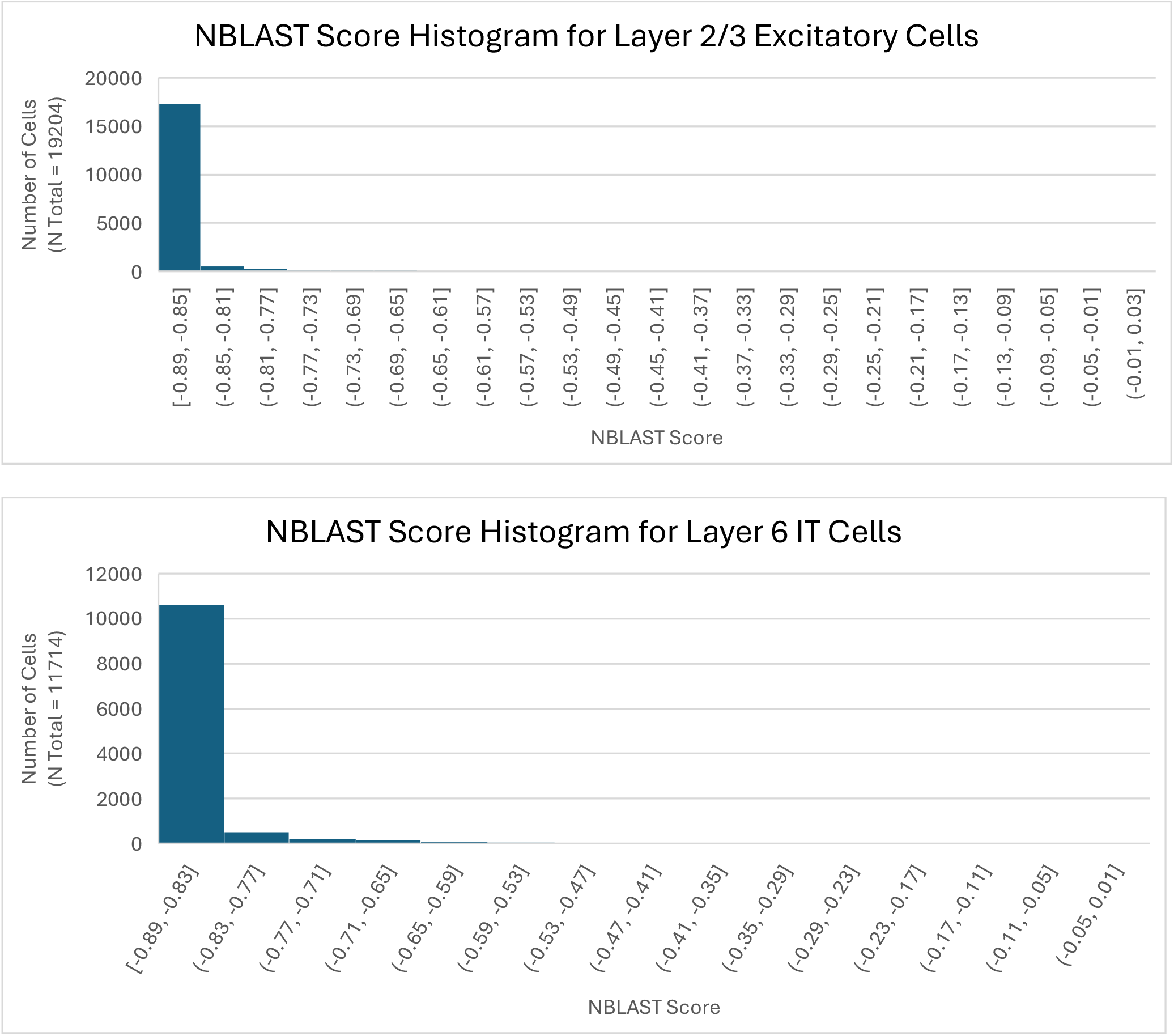
Histogram plots showing the distribution of NBLAST scores for different cell types. As shown in Table 1 below, the mean score for these cells ranged from -0.88 to -0.85. In every case, the threshold NBLAST score of “0” was more than 8 standard deviations from the mean, and in one case 20. Thus, the property of structural similarity as defined by NBLAST analysis is relatively rare for the cells included in this study and does not arise from these cells being of the same class (i.e. IT, ET, NP, etc.)

**Figure 2:**
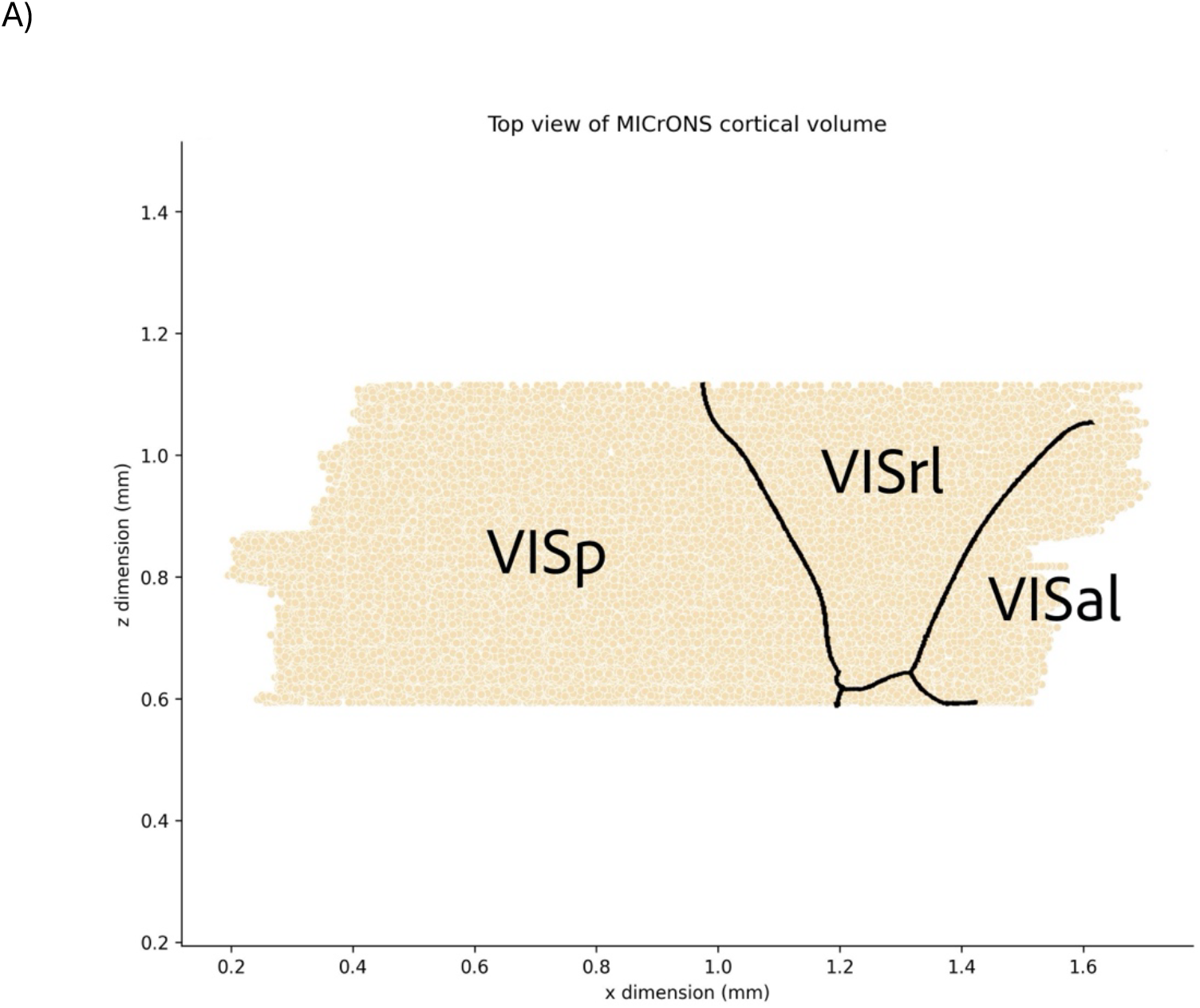
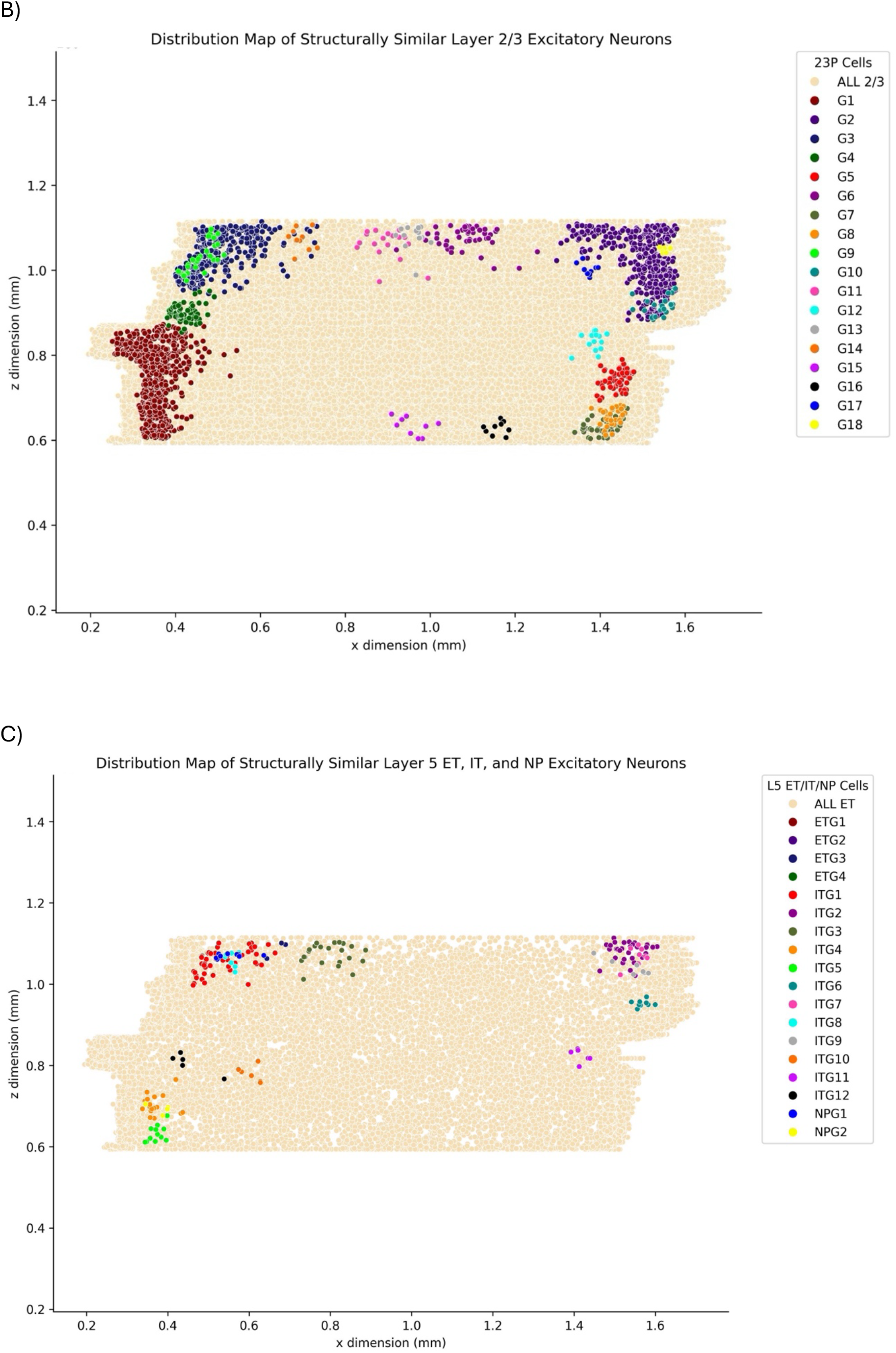
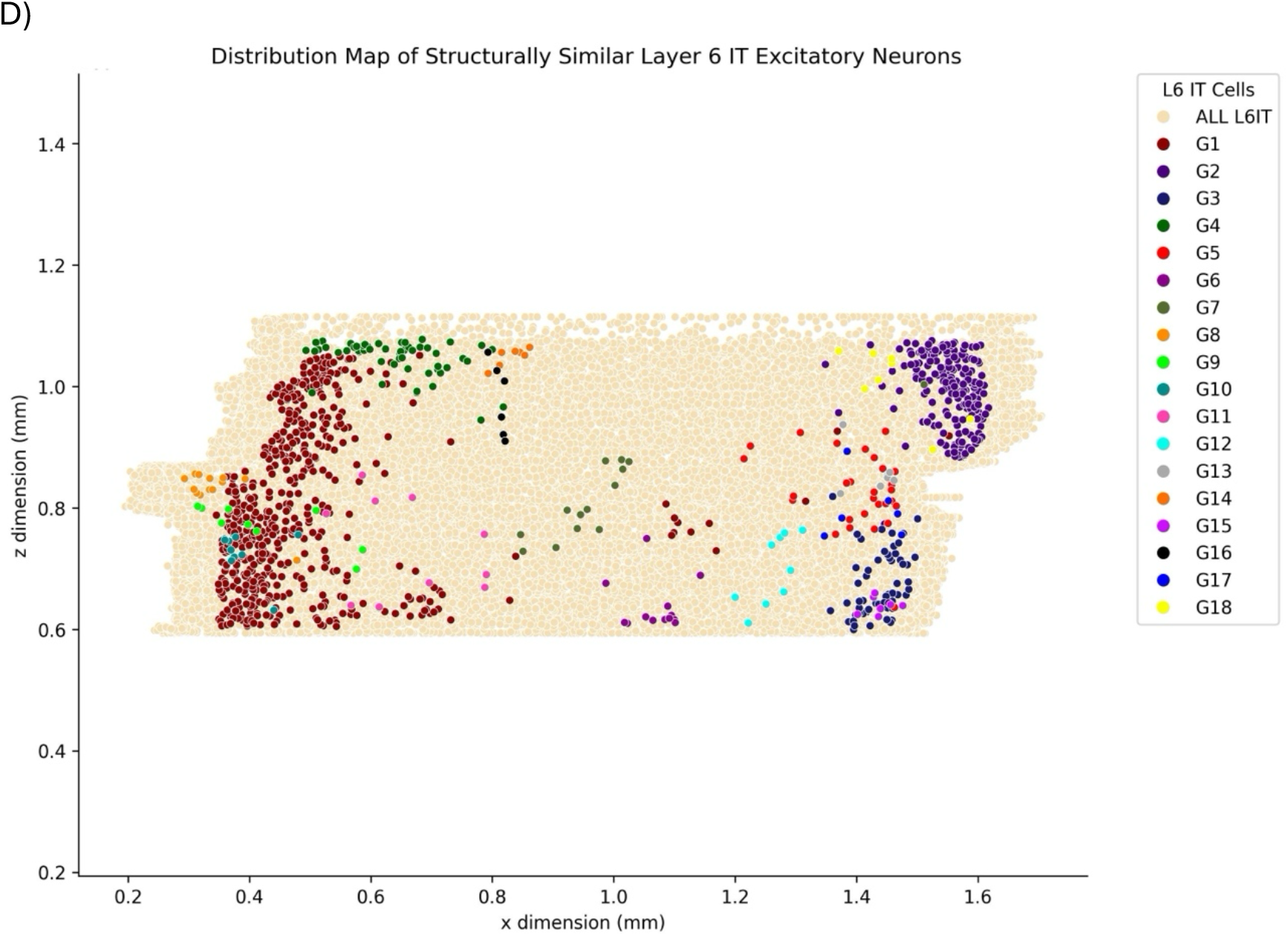
The panels show distribution maps of cell groups defined by structural similarity. The coordinates of cell nuclei were used to position cells. The x-dimension in the volume corresponds most closely to the medial-lateral axis, the z-dimension corresponds most closely to the anterior-posterior axis of the mouse brain. Depth (which is not shown) corresponds to the y-dimension in this dataset. A) shows the tissue outlined by cells from this tissue. The described cortical boundaries for visual areas are also shown (The MICrONS Consortium et al. 2025). B) Layer 2/3 excitatory cells are shown as the background with groups of cells (G1, G2,…) showing significant similarity indicated by group/color. C) Layer 5 ET cells are shown as the background with layer 5 IT, ET, or NP cells showing significant similarity indicated by group/color. D) Layer 6 IT cells are shown as the background with cell clusters showing significant similarity indicated by group/color.

**Table 1:** Summary of Mean NBLAST Scores and Standard Deviatons For the Individual Cell Types.

| Cell Type | Mean Score | Standard Deviation |
| --- | --- | --- |
| L5 ET Cells | -0.88 | +/- 0.04 |
| L5 IT Cells | -0.85 | +/- 0.10 |
| L5 NP Cells | -0.85 | +/- 0.10 |
| L2/3 Excitatory Cells | -0.86 | +/- 0.09 |
| L6 IT Cells | -0.86 | +/- 0.06 |

Within this broader framework, this study set out to test a prediction arising from the Quantized Visual Awareness (QVA) hypothesis: that subsets of primary visual cortex neurons should exhibit unexpectedly strong structural similarity and that these similarities should cluster spatially rather than appear uniformly throughout the cortex (Escobar, 2016, 2013, 2011). QVA postulates that recurrent activation of the primary visual cortex—long known to be essential for generating visual awareness in mammals (Boehler et al. 2008); (Cowey and Walsh 2000); (Kosslyn et al. 2001); (Mehta and Schroeder 2000); (Overgaard et al. 2004); (Pascual-Leone and Walsh 2001); (Silvanto et al. 2005); (Tong 2003)—works in part by repeatedly activating electromagnetic field (EMF) topologies generated by local microcircuits. These EMFs are hypothesized to serve as the physical correlates of elementary perceptual “quanta,” or qualia. Because EMF shape is influenced by both a neuron’s three-dimensional morphology and the spatiotemporal pattern of synaptic input it receives, QVA predicts that neurons capable of generating similar EMF topologies should share partially stereotyped morphological features. Furthermore, these cells should be spatially clustered, analogous to the well-known clustering of orientation-tuned cells into pinwheels (Bartfeld and Grinvald 1992); (Montcastle 1997); (Crair et al. 1997) or color-tuned cells into blobs (Hubel and Livingstone 1983); (Livingstone and Hubel 1984).

The current structural analysis supports several aspects of this prediction. We find that certain pyramidal neuron populations, including layer 5 ET, IT, and near-projecting cells, show pairwise morphological similarity far exceeding that expected by chance—on the order of 8–10 standard deviations above the dataset’s empirical distribution (Figure 1). These similarities are rare when considered across the full population of reconstructed neurons, making the observed degree of convergence surprising under current models of cortical development or connectivity. Moreover, these structurally similar neurons are not randomly interspersed throughout VISp. Instead, they form spatial clusters that include multiple excitatory cell classes, suggesting an underlying organizational principle not captured by existing descriptions of mouse visual-cortical micro-architecture.

One potential alternative explanation is that these similarities arise from lineage relationships, with clonally related neurons inheriting similar morphologies from a common progenitor. However, such a mechanism would be expected to produce clusters distributed broadly and consistently across the cortical sheet. The sparse and spatially selective clustering observed here does not align with that prediction. Instead, the clustering pattern more closely resembles the patch-like organization of known functional maps—such as orientation domains, color-biased patches—suggesting that local circuit demands, rather than developmental lineage alone, may shape these recurring morphologies.

Another possible interpretation is that these cell-topology groups are just another marker for well-known structures like ocular dominance columns. This, however, would not explain why there are so many distinct groupings. What we see is a high level of structural similarity within each cell-topology group, which coincidentally is a relatively rare characteristic (when considering the total number of cells of a certain type) and cell-topology groups that are distinct from each other. In addition, these clusters of cells tend to localize in the same areas of cortical tissue. Taken together, all of these groups may be identifying a row of ocular dominance columns, but at the level of individual cell-topology groups, there must be another phenomenon at play.

A potential concern regarding this work is that the observed morphological similarities could arise as an artifact of tissue boundaries or truncation of neuronal processes within the reconstructed volume. Although many of the identified clusters are located near the lateral edges of the dataset, the spatial distribution of these groups is not consistent with a simple truncation-based explanation. If structural similarity were driven primarily by incomplete reconstructions at tissue boundaries, neurons sharing similar truncation patterns would be expected to appear broadly along multiple edges of the volume rather than as discrete, localized assemblies. Instead, the identified groups form spatially coherent clusters restricted to specific cortical regions, and several cluster members are positioned substantial distances from tissue boundaries. The selective and localized nature of these distributions therefore argues against truncation as the primary source of the observed morphological convergence.

The distribution of cell clusters varies throughout the cortical layers. In layers 2 and 3, there is robust clustering along the “sides” of the tissue with many groups containing large numbers of cells (Figure 2B). In layer 5 the clustering is sparse, with fewer clusters, each containing fewer cells. Nevertheless, the neural cells still collocate into tight groups (Figure 2C). In layer 6, the clustering again is abundant (larger number of groups with a greater number of cells) but more diffuse than the other layers (Figure 2D). Importantly, cell clusters share the same general medial-lateral and anterior-posterior coordinates in each layer, indicating these clusters sit on top of each other, possibly in a columnar fashion.

Extensive studies of the primate primary visual cortex (V1) have uncovered highly organized local structures called ocular dominance columns, each of which corresponds to a point of the visual field (Figure 3). Within each ocular dominance column, exist 50 – 80 microcolumns (also known as minicolumns) defined by the bundle of apical dendrites that arise from cells in layers 2, 3 and 5 and reach up to layer 1 (Von Bonin and Mehler 1971); (Peters and Sethares 1991). Recent evidence indicates that the mouse primary visual cortex is structured in a similar fashion that includes ocular dominance columns (Takahata 2024); (Goltstein 2025).

**Figure 3:**
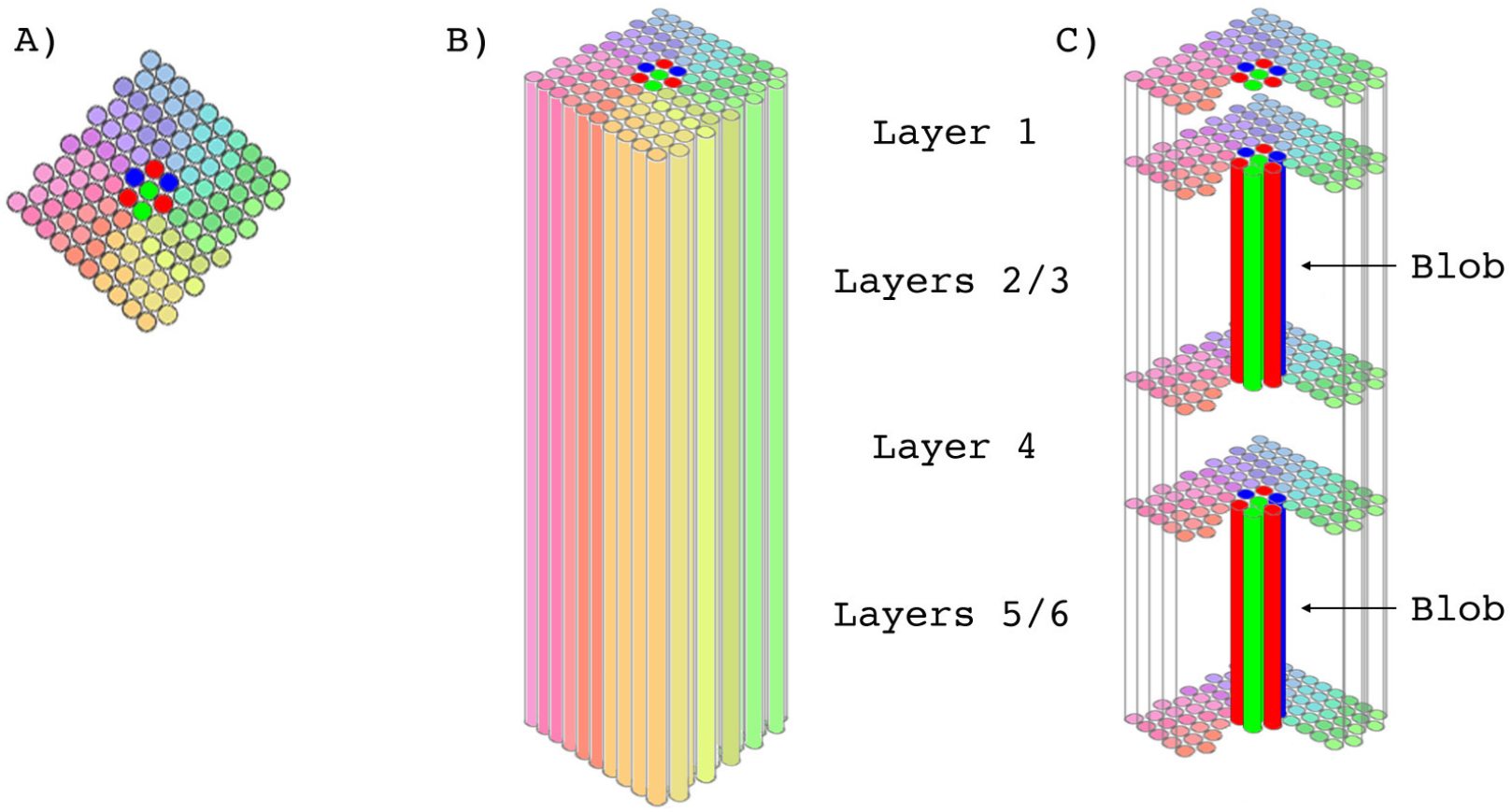
Adapted from Escobar, 2016. A) Top of an ocular dominance column (ODC). The circles are the tops of the microcolumns contained within an ODC as shown in B). The various hues correspond to orientation tuning covering a complete set of 180 degrees. The red, blue and green microcolumns are used to process color. C) These columns pass through blobs – tissue that stains for cytochrome oxidase activity and is tuned to color stimuli.

QVA proposes that the recurrent activation of ocular dominance columns is the mechanism by which individual circuits producing qualia are recruited into a phenomenal visual experience. Although it is not clear from the results of this study, the overlapping architecture observed for cell clusters in the mouse visual cortex is consistent with the idea that individual cells of clusters found at various depths (layers) are located within the same ocular dominance columns. If so, this would provide a mechanism by which cluster cells found in the different layers could all be recruited together as a single unit.

Within the framework of QVA, these observations suggest a plausible anatomical substrate for the generation of elementary EMF topologies. A substantial fraction of the identified neurons—particularly layer 5 extratelencephalic (ET) pyramidal cells, characterized by expansive apical tufts and strong engagement in recurrent feedback from higher visual areas—are well positioned to integrate recurrent inputs terminating in layer 1 and to support large, spatially structured dendritic current flows. In this view, stereotyped EMF configurations could emerge from the interaction between conserved features of single-cell morphology and patterned synaptic drive arising from metastable attractor states in network activity space, as proposed in prior theoretical models. Whether these fields contribute directly to perceptual experience remains an open question, but the presence of repeated morphological motifs provides an empirical foothold for exploring how structure and functional feedback might interact at the level of local microcircuits.

Clearly, these results go beyond what was initially predicted by the QVA hypothesis. The property of cell structure similarity appears to be more widespread and is not only found in the excitatory cells of layer 5 but also in Layers 2, 3, and 6. In addition, clusters of cells displaying cell structure similarity are found in VISp and in the visual rostrolateral and anteriolateral cortex. It may be possible all of these areas are playing a role in generating bits of visual consciousness. The QVA hypothesis will need to be modified to consider what is being observed here.

More broadly, this work highlights how dense connectomic datasets can uncover mesoscale organizational principles that are not apparent from physiology alone. If the visual cortex indeed contains repeating structural motifs that align with qualitative “building blocks” of visual awareness, this would parallel a common theme in biological systems: complex functions emerging from the coordinated activity of many simple, repeated units. Establishing whether these motifs correspond to functional microcircuits, and whether their EMF dynamics carry perceptual significance, will require integrating structural connectomics, large-scale physiology, and computational modeling. The present findings provide a structural foundation on which these next steps can be built.

## Acknowledgements

Special thanks to Bethanny Danskin for her help as part of the Vortex program. Her help was invaluable in learning how to interface with the MICrONS Dataset. I believe the Vortex Program serves as an excellent model for any large connectomics project that is publicly available since this type of interface between the scientific community and the database greatly facilitates its use by the research community.

